# *Photoperiod decoder 1* regulates seasonal changes in energy metabolism through the growth hormone signaling pathway

**DOI:** 10.1101/2025.02.19.638565

**Authors:** Tomoya Nakayama, Taiki Yamaguchi, Michiyo Maruyama, Satoshi Ansai, Makoto Kashima, Romain Fontaine, Christiaan Henkel, Kiyoshi Naruse, Takashi Yoshimura

## Abstract

Seasonal changes in metabolism are crucial for animals to adapt to annual environmental fluctuations. However, the molecular mechanisms underlying these adaptations remain poorly understood. Here, we identified a novel gene, *photoperiod decoder 1* (*phod1*), which exhibits a unique bimodal expression pattern under long-day conditions in Japanese medaka fish (*Oryzias latipes*). While *phod1* is conserved across many vertebrates, except for eutherians, its function remained unknown. Single-cell RNA sequencing analysis revealed that *phod1* is predominantly expressed in specific pituitary cells that co-express opsin and circadian clock genes. Transcriptomic analysis using *phod1* knockout fish demonstrated that *phod1* is essential for photoperiodic regulation of growth hormone. Furthermore, transcriptomic and metabolomic analyses of the liver, the primary target of growth hormone, revealed significant alterations in energy metabolism. Behavioral analysis also showed that *phod1* knockout fish exhibited significantly reduced locomotor activity. These findings indicate that *phod1* plays a crucial role in seasonal metabolic adaptation through modulation of the growth hormone signaling pathway.

---

Metabolic regulation is a fundamental aspect of life that is crucial for maintaining homeostasis and adapting to environmental changes. In many organisms, particularly those living in temperate regions, the metabolism undergoes significant seasonal fluctuations. For example, many animals coordinate metabolism with reproduction and energy allocation, with extensive remodeling of energy storage and utilization to ensure the optimal timing of reproduction and survival (Parker and Cheung, 2020). Migratory species undergo dramatic physiological changes as breeding season approaches, including increased metabolic rates, hypertrophy of the locomotor muscles and heart, and enhanced lipid metabolism to support their energetically costly seasonal movements (Ramenofsky and Wingfield, 2007). Hibernating animals dramatically lower their metabolic rates during winter to overcome harsh environmental conditions (Klug and Brigham, 2015). These examples not only illustrate the diverse ways in which animals adapt their metabolism to seasonal changes, but also emphasize the significance of the synchronization of metabolism with seasonal environmental changes.

The ability to anticipate and prepare for these seasonal changes through adjustments in physiological functions and behaviors is a key process for animals living in temperate regions. Central to this adaptive mechanism is photoperiodism, which is the ability of organisms to detect and respond to changes in day length (Rowan, 1925; Follett and Sharp, 1969). Because photoperiodic changes are repeated at a fixed time each year, they serve as reliable environmental cues for upcoming seasonal changes, allowing animals to initiate physiological and behavioral adjustments well in advance of actual environmental shifts. This remarkable system enables organisms to synchronize their various biological processes with the annual cycle, ensuring optimal timing for energetically demanding activities, such as breeding, migration, or dormancy (Bradshaw and Holzapfel, 2007). Recent advances in molecular biology have significantly enhanced our understanding of the core pathway of the photoperiodic regulation of reproduction in vertebrates (Yoshimura et al., 2003; Nakao et al., 2008; Ono et al., 2008; Nakane et al., 2013; Wood et al., 2015; Wood et al., 2020). While the molecular mechanisms underlying seasonal reproduction are well understood, our knowledge of seasonal metabolic regulation remains limited, particularly in non-mammalian vertebrates. Additionally, although photoperiodic regulation of metabolic processes has been reported in various vertebrates (Fish: Biswas and Takeuchi, 2002; Bird: Boon et al., 2000; Mammal: Mariné-Casadó et al., 2018; Small et al., 2023), the pathways linking photoperiodic sensing to metabolic shifts are not well understood.

Here, we identified a novel gene that displays distinct photoperiodic expression patterns using Japanese medaka (*Oryzias latipes*), an excellent model organism for seasonal adaptation studies due to its highly sophisticated seasonal responses (Egami, 1954; Awaji and Hanyu, 1989; Shimmura et al., 2017; Nakayama et al., 2019; Nakayama et al., 2020; Nakayama et al., 2023) and extensive genomic resources, including accurate genomic information (Kasahara et al., 2007; Ichikawa et al., 2017) and well-established genome-editing tools (Ansai and Kinoshita, 2014). This gene, which we named *photoperiod decoder 1* (*phod1*), exhibited a unique bimodal expression with peaks in the morning and evening, specifically under long-day conditions. *phod1* is conserved across many vertebrates, but is absent in most mammals, suggesting a potentially important role in seasonal adaptation in non-mammalian vertebrates. However, the functions of *phod1* were not previously characterized in any vertebrates. In this study, we found that *phod1* is essential for photoperiodic regulation of growth hormone signaling and plays a crucial role in seasonal metabolic adaptation.

## Results

### Identification of an uncharacterized gene that shows photoperiodic responsiveness

Previously, we conducted genome-wide expression analysis in the brain region containing the hypothalamus and the pituitary of female medaka during the transition from short-day (10 h light / 14 h dark; 26°C) to long-day (16 h light / 8 h dark; 26°C) conditions, identifying 1,249 photoperiodically regulated transcripts (Nakayama et al., 2019). Further analysis of this dataset revealed a transcript, *LOC101165601*, that exhibited a distinctive bimodal expression pattern, with peaks in the morning and around dusk under long-day conditions (Fig. 1a; Supplementary Fig. 1a). This characteristic expression profile was not observed under short-day conditions but was observed on the first and second days after the transition to long-day conditions. According to the National Center for Biotechnology Information (NCBI) database, *LOC101165601* was reported as an uncharacterized protein. Previously generated ribosome profiling data also supported the translation of this gene into a protein (Supplementary Fig. 1b) (Nakayama et al., 2019), although no known protein domains were identified. Ortholog analysis using ORTHOSCOPE (Inoue and Satoh, 2019) indicated that *LOC101165601* is conserved among fish, amphibians, reptiles, birds and marsupials but was not detected in eutherians (Supplementary Fig. 2). Notably, the function of genes that appeared to be orthologs in any vertebrates is unknown. Therefore, we conducted a more detailed analysis of *LOC101165601*. We next examined the expression profile of *LOC101165601* in other tissues under short- and long-day conditions. Although we found *LOC101165601* to be expressed in the brain and pituitary gland, as well as in the gills, bimodal expression was observed only under long-day conditions in the brain and pituitary gland (Supplementary Fig. 3). Long-day-specific bimodal expression of *LOC101165601* in the brain and pituitary gland was also observed in the male, suggesting that there are no sex differences in the regulation of *LOC101165601* expression (Supplementary Fig. 4). To further characterize the expression profile of *LOC101165601*, we maintained medaka under long-day conditions and transferred them to short-day conditions. Consistent with the above results, a bimodal expression with peaks in the morning and evening was observed under long-day conditions, but not after the transition to the short-day conditions (Fig. 1b). The expression profile of *LOC101165601* was also examined in medaka transferred from long-day conditions to continuous light (LL) or dark (DD) conditions. After the transition to LL conditions, *LOC101165601* expression was found to be remained high (Fig. 1c). In contrast, bimodal expression or even expression was not detected after the transition to DD conditions, except at 8 h after the transition to DD conditions (Fig. 1d). To examine whether there is a critical day length for the regulation of *LOC101165601* expression, we kept medaka under various photoperiod conditions. Under the 10 h, 11 h, 12 h and 13 h photoperiod conditions, only one peak was observed in the middle of the day; however, when medaka were kept under the 14 h photoperiod conditions, two expression peaks appeared in the morning and evening (Fig. 1e). These results suggest that the light stimulus is important for the induction of *LOC101165601* and that photoperiod conditions significantly alter its expression profile in the brain and pituitary. Since the expression of *LOC101165601* decodes photoperiodic information, we named this previously unknown gene *<u>pho</u>toperiod <u>d</u>ecoder <u>1</u>*(*phod1*).

**Fig. 1.**
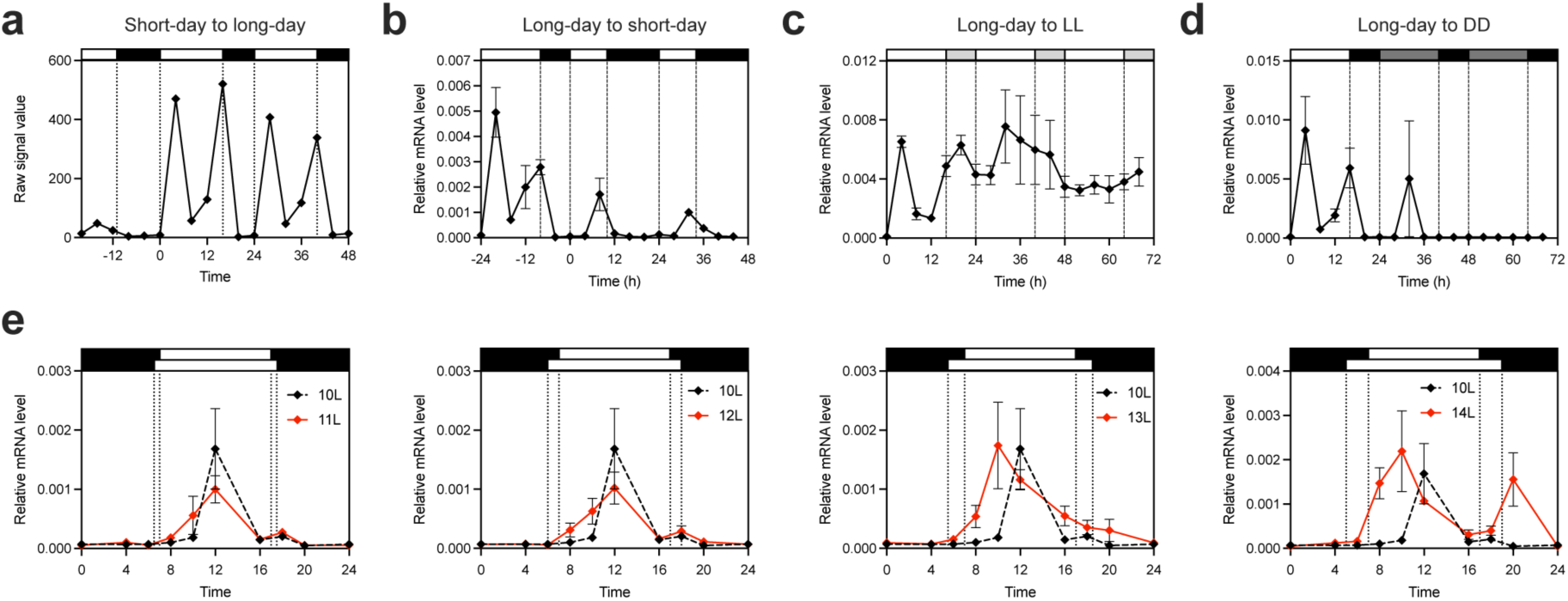
Expression profile of *LOC101165601* under different photoperiod conditions. **a** Expression profile of *LOC101165601* in the brain region containing the hypothalamus and the pituitary of medaka maintained under short-day conditions and transferred to long-day conditions (Nakayama et al., 2019). **b** Expression profile of *LOC101165601* in medaka kept under long-day conditions and transferred to short-day conditions (mean ± SEM, and n = 4). **c** Expression profile of *LOC101165601* in medaka kept under long-day conditions and transferred to constant light conditions (LL) (mean ± SEM, and n = 2 to 5). **d** Expression profile of *LOC101165601* in medaka kept under long-day conditions and transferred to constant dark conditions (DD) (mean ± SEM, and n = 3 to 4). **e** Expression profile of *LOC101165601* in medaka kept under different photoperiod conditions (mean ± SEM, and n = 3 to 4).

### *phod1* is expressed in the pituitary

To examine the expression site of *phod1*, we collected brain and pituitary samples under long-day conditions at ZT4, when *phod1* expression peaked (Fig. 1), and performed in situ hybridization. Interestingly, *phod1* was expressed in the pituitary gland, a key endocrine organ controlling various physiological functions (Fig. 2). Expression signals were mainly found in the proximal pars distalis (PPD) and pars intermedia (PI) and were not clustered together but were scattered in the PPD and PI (Fig. 2b, c). We also detected *phod1* expression in the preoptic area (POA) (Supplementary Fig. 5), although the number of *phod1*-expressing cells was much smaller compared to that in the pituitary gland.

**Fig. 2.**
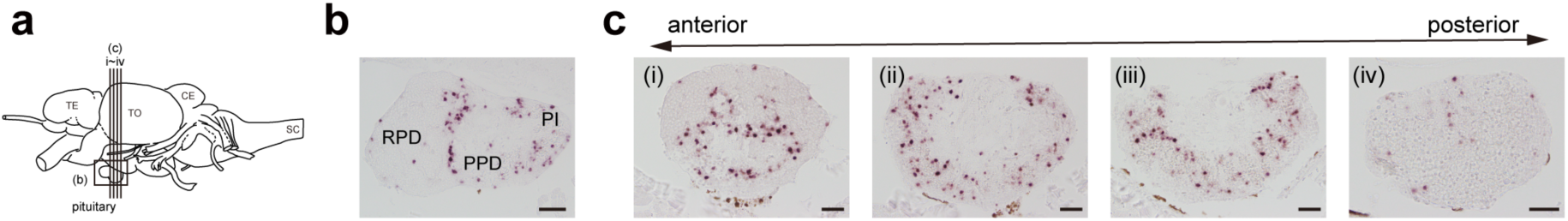
Expression sites of *phod1* in the pituitary gland. **a** Schematic illustration of the medaka brain showing the planes of the sections in b and c. TE, telencephalon; TO, tectum opticum; CE, corpus cerebelli; SC, spinal cord. **b** Spatial distribution of *phod1* expression in sagittal sections of the pituitary gland, visualized by in situ hybridization (scale bar = 50 µm). **c** Spatial distribution of *phod1* expression in coronal sections of the pituitary gland, visualized by in situ hybridization (scale bar = 20 µm).

### *phod1*-expressing cells express opsin genes and clock genes

Given that *phod1* is predominantly expressed in the pituitary, we next analyzed single-cell RNA-seq (scRNA-seq) data from the medaka pituitary deposited in public databases (Siddique et al., 2021) to explore the *phod1* expression in cells in detail. We used the NCBI annotation as a reference because *phod1* is only annotated in the NCBI annotation (note that Ensembl annotation was used in Siddique et al., 2021). Transcriptome information in each cell was obtained from 2,853 cells, and clustering analysis allowed us to identify 15 cell populations (Fig. 3a, left). Following previous annotations, we were able to identify almost the same cell populations as in the previous report (Siddique et al., 2021), including FSH-gonadotropes (*fshb*^+^), LH-gonadotropes 1/2 (*lhb*^+^), thyrotropes (*tshba*^+^), lactotropes 1/2 (*prl*^+^), corticotropes (*pomc*^+^/*tbxt*^+^), melanotropes (*pomc*^+^), somatotropes (*gh1*^+^), somatolactotropes (*somatolactin*^+^), macrophages (*LOC105356008*^+^), and red blood cells (*hemoglobin*^+^). In these cell populations, *phod1* was found to be specifically expressed in the cell population that has been named *glycoprotein hormones, α polypeptide* (*cga*)-expressing cells in a previous report (Siddique et al., 2021) (Fig. 3a, right; Supplementary Fig. 6). In this study, we redefined this cell population as *phod1*^+^ cells. To further characterize *phod1*^+^ cells, we extracted genes that were specifically expressed in *phod1*^+^ cells, and 497 genes were identified as marker genes (fold-change > 5, with adjusted *P* value < 1.0E-50 and expressed in over 80% of *phod1*^+^ cells) (Supplementary Table 1). Among the marker genes, we found genes encoding a photoreceptor [*opsin 5-like 1c* (*opn5l1c*, Gene ID: 101163490)], circadian clock components [*period circadian protein homolog 2* (*per2a*, Gene ID: 101161338), *period circadian regulator 2* (*per2b*, Gene ID: 101158624), *cryptochrome circadian regulator 1b* (*cry1b*, Gene ID: 101160902), *HLF transcription factor, PAR bZIP family member a* (*hlfa*, Gene ID: 101160073)] and endocrine signaling molecules [*inhibin subunit beta Aa* (*inhbaa*, Gene ID: 101170230)] (Fig. 3b). *Somatostatin receptor 2* (*sstr2*, Gene ID: 101172660) and *prolactin releasing hormone receptor 2a* (*prlhr2a*, Gene ID: 101174270) were also contained in the marker genes, which may suggest *phod1*^+^ cells can respond to somatostatin and prolactin releasing hormone signaling. Additionally, we found *GATA binding protein 2* (*gata2*) *b* (*gata2b*, Gene ID: 100125427) was specifically expressed in *phod1*^+^ cells (Fig. 3b). Given that GATA2 is a key transcription factor for the development and maintenance of the gonadotropes and thyrotropes (Scully & Rosenfeld, 2002) and CGA is the common *α* subunit of LH, FSH and TSH, *phod1*^+^ cells may share a common developmental origin with gonadotropes and thyrotropes, as evidenced by the expression of both *gata2a* (Gene ID: 101161391) and *cga* in *phod1*^+^ cells (Fig. 3b). Gene ontology (GO) and Kyoto Encyclopedia of Genes and Genomes (KEGG) pathway enrichment analysis of *phod1*^+^ cell marker genes revealed significant enrichment of terms related to intercellular communications, including “modulation of chemical synaptic transmission,” “regulation of secretion,” and “signaling by receptor tyrosine kinases” (Fig. 3c). These results suggest that *phod1*^+^ cells may play an important role in coordinating intercellular communication within the pituitary. Cell-cell communication analysis using LIANA (Dimitrov et al., 2022) revealed potential ligand-receptor interactions between *phod1*^+^ cells and hormone-producing cell populations (Fig. 3d). The analysis suggested that *phod1*^+^ cells may communicate with other cell populations through multiple ligands, including *inhbaa*, *rln3*, *nrg3b*, and *ncam1* (Supplementary Fig. 6). Specifically, these cells showed interactions with LH-gonadotropes through *inhbaa*, FSH-gonadotropes through *inhbaa* and *nrg3b*, somatotropes through *inhbaa* and *rln3*, somatolactotropes through *inhbaa* and *ncam1*, corticotropes through *rln3* and lactotropes through *nrg3b*, suggesting that *phod1*^+^ cells might serve as important mediators of intercellular communication within the pituitary.

**Fig. 3.**
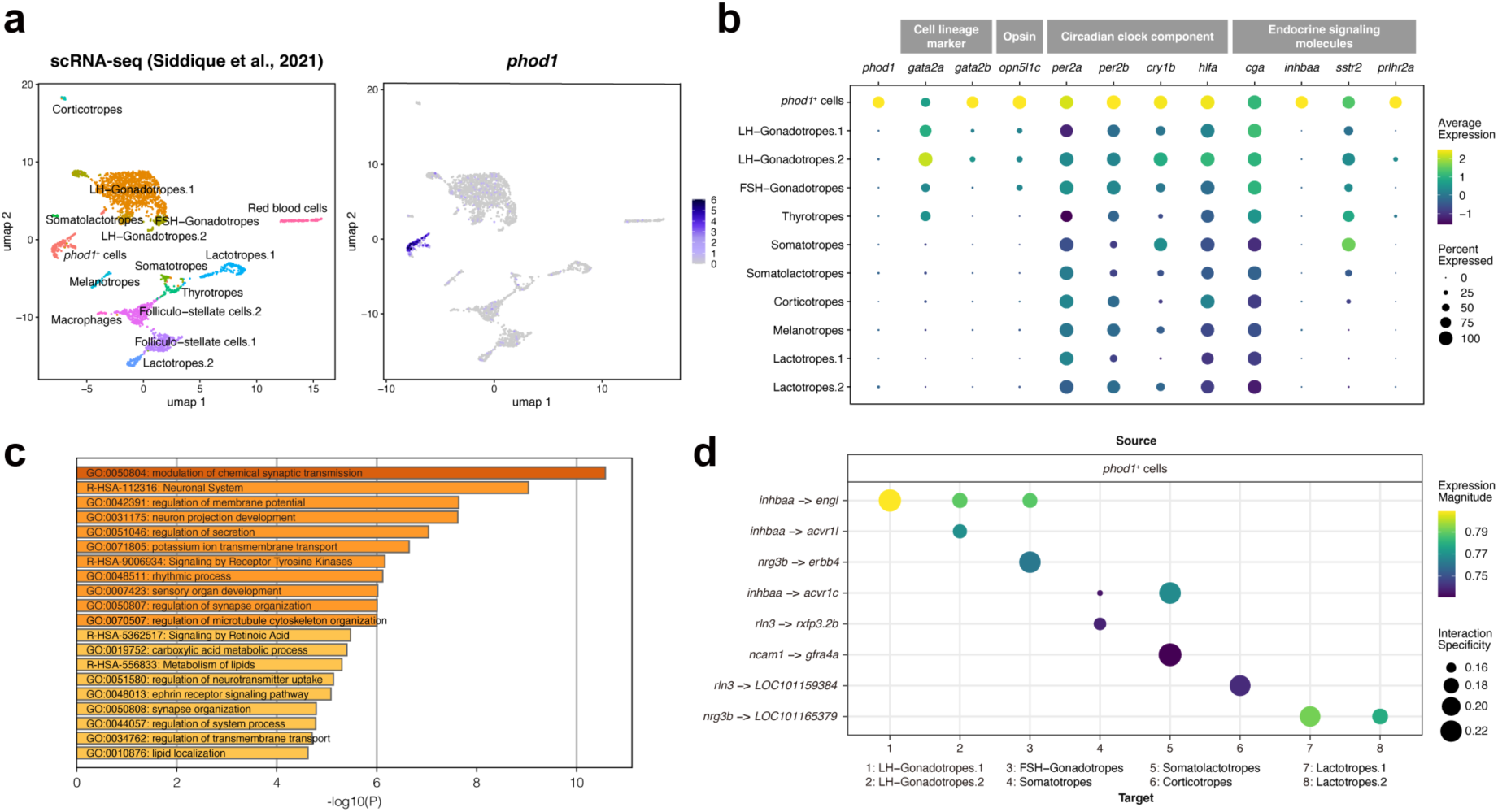
Characterization of *phod1*-expressing cells in the pituitary gland using single-cell RNA sequencing. **a** Left: Uniform Manifold Approximation and Projection (UMAP) plot of pituitary cell populations colored according to cell type. Right: Expression level of *phod1* projected onto the UMAP plot. Each dot represents a single cell. **b** Expression patterns of cell type-specific marker genes across the identified cell populations. Dot size and color intensity indicate the percentage of cells expressing each gene and the average expression level, respectively. **c** GO and KEGG pathway enrichment analysis of 497 genes identified as marker genes in *phod1*^+^ cells. **d** Dot plot of cell-cell communication patterns from *phod1*^+^ cells to hormone-producing cell populations. The y-axis shows ligand-receptor pairs, and the x-axis shows target cell populations. Dot size and color intensity indicate the interaction specificity and the expression magnitude, respectively.

### Disruption of *phod1* affects photoperiodic regulation of growth hormone

To understand the physiological and ecological functions of *phod1*, we next generated *phod1* knockout medaka using the CRISPR/Cas9 system. The genomic region containing the transcription and translation start site of *phod1* was deleted using two different guide RNAs (Fig. 4a, b). We crossed an injected G_0_ founder with a wild-type fish to obtain F_1_ heterozygotes (*phod1*^+/−^) and then crossed the F_1_ heterozygotes to generate F_2_ knockout (*phod1*^−/−^) and wild-type (*phod1*^+/+^) siblings. Expression of *phod1* was significantly abolished in the knockout medaka, indicating this deletion efficiently knocked out the *phod1* expression (Fig. 4c). To examine the effects of *phod1* knockout on the expression profiles, we performed RNA-sequencing (RNA-seq) analysis using the brain region containing the hypothalamus and the pituitary collected (sampled at the midpoint of the light period, 12:00) from adult medaka of each genotype before and 12 days after the transition from short-day to long-day conditions. First, we compared the transcriptome between short- and long-day conditions in wild-type to extract genes regulated by photoperiod. In this analysis, 111 genes including several hormone genes (*follicle-stimulating hormone β* [*fshb*], *luteinizing hormone subunit β* [*lhb*], *cga*, and *growth hormone 1* [*gh1*]) were identified as photoperiodically regulated genes (adjusted *P* value < 0.01) (Supplementary Table 2). Next, we compared the transcriptomes of wild-type and knockout medaka under short- and long-day conditions and identified 207 and 262 differentially expressed genes (DEGs), respectively (Fig. 4d and Supplementary Tables 3, 4). GO and KEGG pathway enrichment analysis of DEGs in long-day conditions highlighted enrichment of genes related to “Cholesterol biosynthesis,” “Intra-Golgi and retrograde Golgi-to-ER traffic,” “regulation of lipid metabolic process,” “neural crest cell development,” “cellular response to nutrient levels” and “TCA cycle” (Fig. 4e). A Venn diagram of these DEGs and photoperiodically regulated genes showed that 13 genes overlapped (Fig. 4f). Notably, two hormone genes that are expressed in the pituitary were included among these common genes: *gh1* and *fshb* (Fig. 4f, Supplementary Fig. 7a). While *gh1* expression was induced by a long-day stimulus in wild-type fish (Supplementary Fig. 8), *phod1* knockout fish exhibited constitutively high *gh1* expression regardless of the photoperiod (Fig. 4g). Because the expression level of *gh1* could be changed by the photoperiod, we hypothesized that the duration of exposure to short- or long-day conditions might affect its expression levels. Therefore, we examined *gh1* expression in medaka that were maintained under long-day conditions and then transferred them to short-day conditions, comparing the expression levels before and 14 days after the transition. As a result, we found the relative expression levels of *gh1* between wild-type and *phod1* knockout fish showed opposite patterns: higher expression in *phod1* knockout fish in the experiment from short-day to long-day conditions (Fig. 4g), while lower expression in *phod1* knockout fish in the experiment from long-day to short-day condition (Fig. 4h). In addition, whereas wild-type fish showed a clear reduction in *gh1* expression in response to short-day stimulus, *phod1* knockout fish showed no such photoperiodic response. These results suggested that *phod1* plays a crucial role in mediating the photoperiodic regulation of *gh1* expression. Since *fshb* is known to be important for gonadal development in fish (Levavi-Sivan et al., 2010) and the expression of *fshb* was downregulated in *phod1* knockout fish under long-day conditions (Supplementary Fig. 7a), we examined the effect of *phod1* knockout on gonadal development. Although *phod1* knockout female fish tended to have a lower gonadosomatic index (GSI), we found no statistically significant differences in the degree of gonadal development between the wild-type and knockout fish (Supplementary Fig. 7b). Therefore, we focused on subsequent analyses of the effects of growth hormone.

**Fig. 4.**
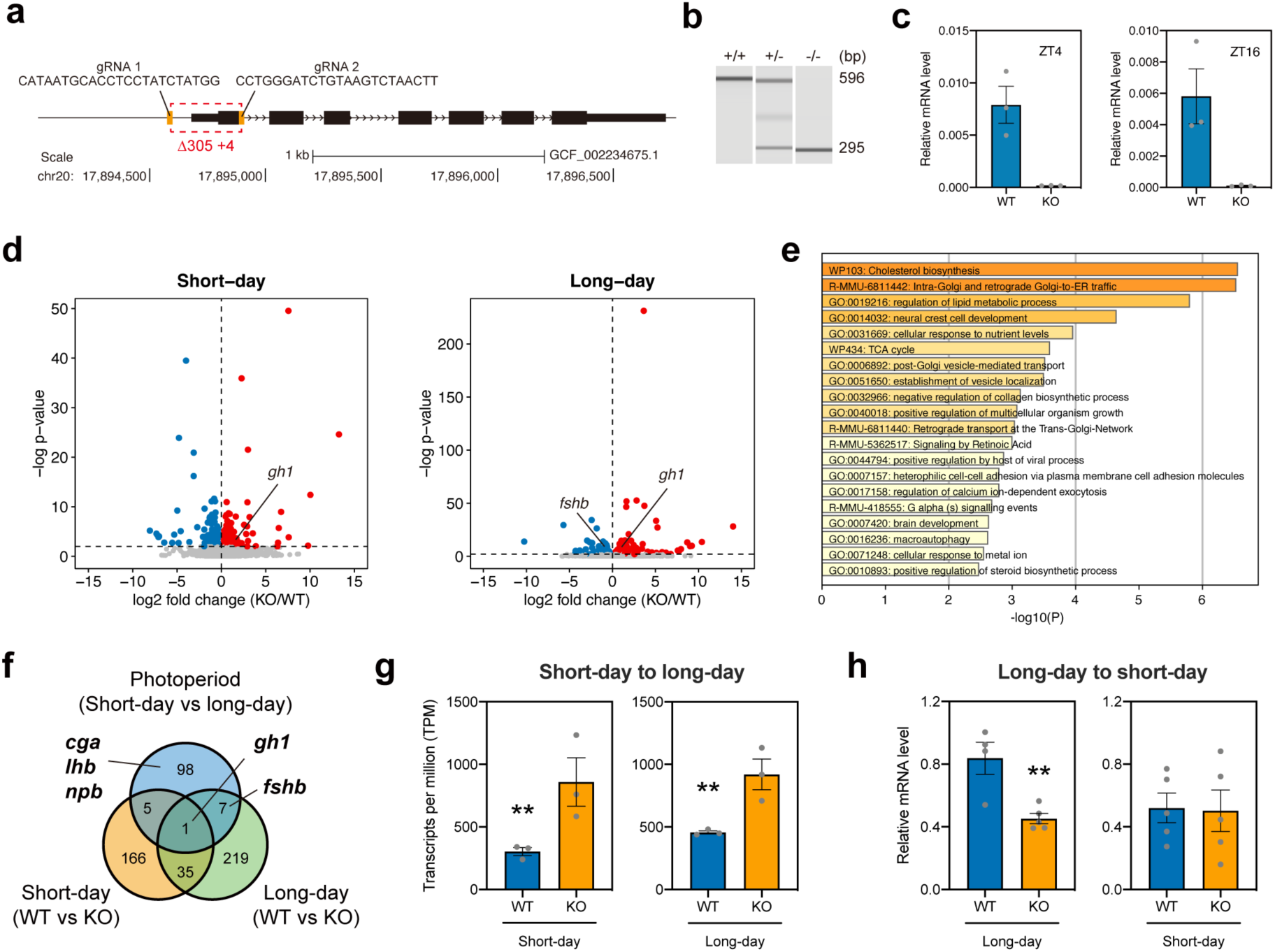
Transcriptional changes induced by *phod1* knockout in the brain. **a** Genomic region targeted and deleted by CRISPR/Cas9 guide RNAs. Black bars represent *phod1* exons. The target sequences of the guide RNAs (gRNA) are shown above the deleted region (red dotted lines). **b** PCR fragments obtained from the genomic DNA of wild-type (596 bp), heterozygous mutant (596 and 295 bp), and knockout (295 bp) fish were analyzed using a microchip electrophoresis system. **c** Expression levels of *phod1* measured by qPCR in wild-type and knockout medaka kept under long-day conditions. The left panel shows the expression level at ZT4 when the morning peak was observed. The right panel shows the expression level at ZT16 when the evening peak was observed. Each dot represents an individual dataset. **d** Volcano plot of differentially expressed genes (DEGs) for each comparison. Red, blue, and gray dots indicate upregulated, downregulated, and unchanged genes, respectively (adjusted *P* value < 0.01). **e** GO and KEGG pathway enrichment analysis of DEGs under long-day conditions. **f** Venn diagram showing the overlap of each DEGs. **g** Expression levels of *gh1* in wild-type and knockout medaka kept under short-day conditions and transferred to long-day conditions. The expression levels under long-day conditions are data from the 12^th^ day after the transition from short-day to long-day conditions (DESeq2 Wald test, **adjusted *P* value < 0.01, mean ± SEM, and n = 3). **h** Expression level of *gh1* in wild-type and knockout medaka kept under long-day and transferred to short-day conditions (Student’s t-test, \*\**P* < 0.01, mean ± SEM, and n = 4 to 5). The expression levels under short-day conditions were measured from the 14^th^ day after the transition from long-day to short-day conditions.

### Loss of *phod1* leads to altered hepatic metabolism and reduced locomotor activity

To explore the downstream effects of *phod1* in the peripheral tissue, we conducted RNA-seq analysis on the liver, the primary target of growth hormones, and compared wild-type and knockout medaka under long-day conditions. This analysis revealed 355 DEGs (adjusted *P* value < 0.01) (Fig. 5a and Supplementary Table 5). GO and KEGG pathway enrichment analyses of these DEGs revealed several significantly enriched biological processes (Fig. 5b), including “carbohydrate metabolic process,” “monocarboxylic acid metabolic process,” “small molecule biosynthetic process,” “amino acid metabolic process,” and “protein processing in endoplasmic reticulum”. These findings suggest that the DEGs are involved in diverse metabolic pathways and cellular processes. Notably, genes related to the “regulation of insulin-like growth factor (IGF) transport and uptake by insulin-like growth factor binding proteins (IGFBPs) ” were also enriched, including *igf2* and *igfbp2*, which showed altered expression in *phod1* knockout fish (Fig. 5c), indicating a potential impact on the growth hormone signaling pathway. In addition, the expression of several key metabolic genes was changed in *phod1* knockout fish, including those involved in carbohydrate metabolism (*glucokinase* [*gck*], *glucose-6-phosphatase* [*g6pc*], and *fructose-1,6-bisphosphatase 1* [*fbp1*]) (Bian et al., 2022), lipid metabolism (*proliferator-activated receptor alpha* [*ppara*], *carnitine O-palmitoyltransferase 1* [*cpt1a*], and *fatty acid synthase* [*fasn*]) (McGarry and Brown, 1997; Liu et al., 2012; Wang et al., 2015), and amino acid metabolism (*glutamate dehydrogenase* [*glud1*] and *aldehyde dehydrogenase 6 family member A1* [*aldh6a1*]) (Fig. 5c). To further examine the metabolic consequences of *phod1* knockout, we performed a metabolomic analysis of livers from wild-type and knockout medaka and found 15 metabolites to be significantly different between wild-type and knockout fish (Supplementary Fig. 9). Notably, there were significant changes in various metabolites involved in cellular energy metabolism (Fig. 5d). These included metabolites involved in glycolysis and the pentose phosphate pathway (glucose 6-phosphate, fructose 6-phosphate, and ribulose 5-phosphate), coenzymes and cofactor (NAD, NADPH, FAD, FMN, and CoA), a nucleotide (ADP), nucleosides (guanosine and inosine), and amino acids (arginine and asparagine) (Fig. 5d) (Supplementary Fig. 9). The accumulation of multiple coenzymes and metabolic intermediates suggests that *phod1* knockout leads to significant perturbations in energy metabolism, particularly in the efficiency of metabolite utilization in key cellular processes. Finally, to assess how the metabolic changes induced by *phod1* deletion affected whole-body activity, we conducted a behavioral analysis comparing wild-type and *phod1* knockout medaka. Medaka maintained under short-day conditions were transferred to long-day conditions, and their locomotor activity was tracked during the transition. As a result, we found that while wild-type fish gradually increased their locomotor activity in response to long-day stimulus, *phod1* knockout fish exhibited consistently low activity levels (Fig. 5e). Quantification of the total locomotor activity during both light and dark phases revealed that knockout fish showed significantly reduced activity regardless of lighting conditions (Fig. 5f). Taken together, our integrated analyses combining transcriptomics, metabolomics, and behavioral assessments revealed that *phod1* knockout results in extensive remodeling of hepatic metabolism and reduced physical activity. These findings highlighted the critical role of *phod1* in maintaining metabolic homeostasis and behavior to adapting to seasonal environmental changes.

**Fig. 5.**
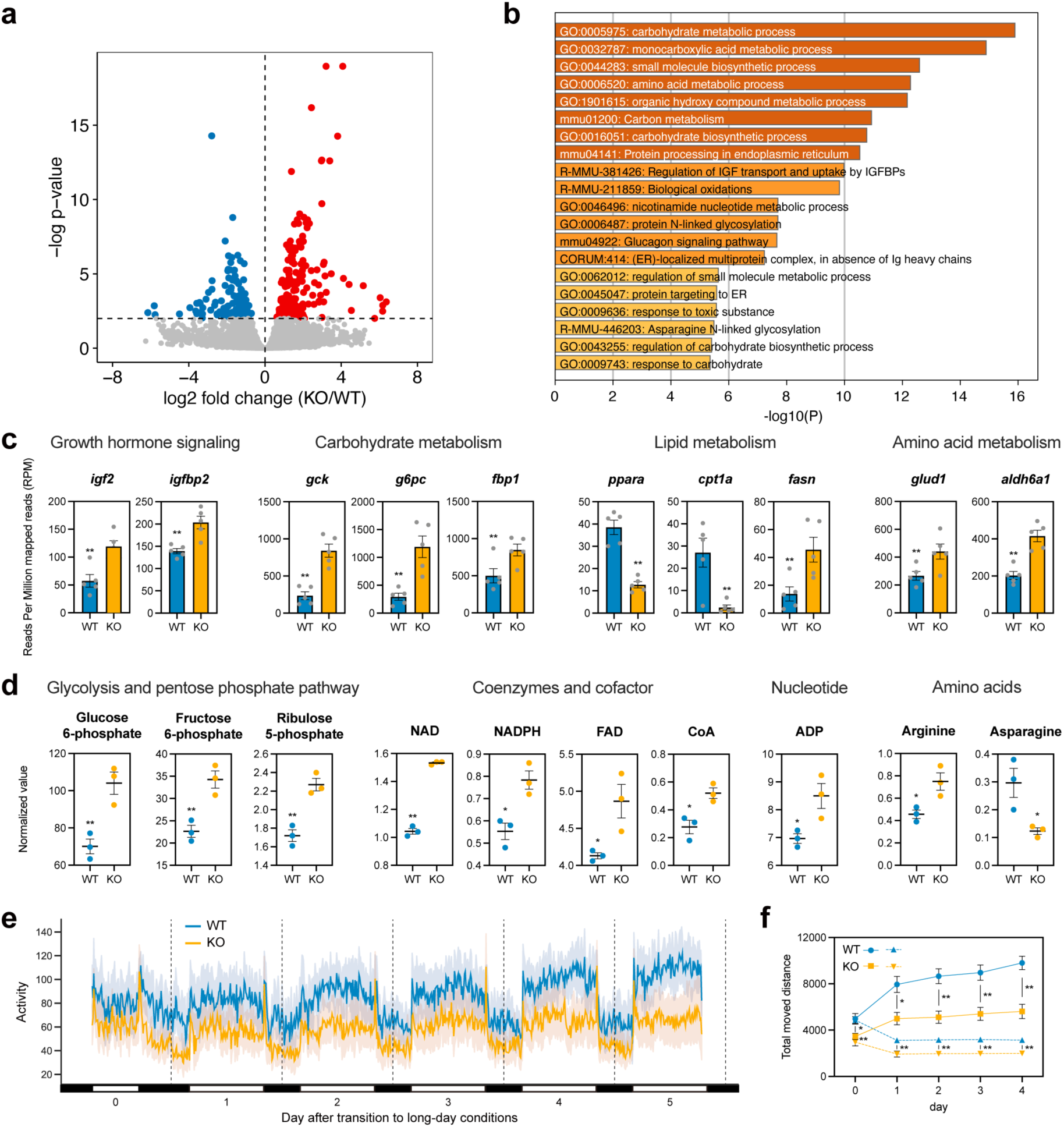
Transcriptomic, metabolomic, and behavioral impacts of *phod1* knockout. **a** Volcano plot of differentially expressed genes (DEGs) in each comparison. Red, blue, and gray dots indicate upregulated, downregulated, and unchanged genes, respectively (adjusted *P* value < 0.01). **b** GO and KEGG pathway enrichment analysis of DEGs in the liver. **c** Expression levels of key genes involved in metabolism in wild-type and knockout fish (DESeq2 Wald test, **adjusted *P* value < 0.01, mean ± SEM, and n = 5). **d** Normalized values of metabolites involved in metabolism in wild-type and knockout fish (Student’s *t*-test, \**P* < 0.05, \*\**P* < 0.01, mean ± SEM, and n = 3). **e** Locomotor activity of wild-type (blue line) and knockout medaka (orange line) after transitioning to long-day conditions. **f** Total locomotor activity of wild-type (blue) and knockout medaka (orange) in both light (solid line) and dark phases (dotted line) (Two-way repeated measures ANOVA: genotype effect, light: F_(1, 26)_ = 17.01, \*\**P* = 0.0003, dark: F_(1, 26)_ = 23.43, \*\**P* < 0.0001; time effect, light: F_(2.488, 64.70)_ = 67.79, \*\**P* < 0.0001, dark: F_(1.638, 42.59)_ = 35.84, \*\**P* < 0.0001; genotype × time interaction, light: F_(4, 104)_ = 9.518, \*\**P* < 0.0001, dark: F_(4, 104)_ = 2.545, *P* = 0.0438; Šídák’s multiple comparisons test, *P* < 0.05 at all time points; mean ± SEM, n = 13 to 15).

## Discussion

In this study, we identified and characterized a novel gene, *photoperiod decoder 1* (*phod1*), in medaka (*Oryzias latipes*). Using a comprehensive approach that combines scRNA-seq, RNA-seq, metabolomic analysis, and genome editing techniques, we demonstrated that *phod1* plays a crucial role in seasonal metabolic adaptation. Our findings revealed that *phod1* exhibits a unique expression pattern in response to photoperiodic changes and influences various aspects of metabolism and physiology, including growth hormone signaling and energy metabolism. Our results provide new insights into the molecular mechanisms underlying seasonal metabolic adaptations in vertebrates.

The bimodal expression profile of *phod1* observed under long-day conditions was tightly regulated by photoperiod, offering intriguing insights into its photoperiodic control. In our experiments, we compared *phod1* expression under short-day (lights on at 7:00, off at 17:00) and long-day (lights on at 4:00, off at 20:00) conditions. A long day stimulus was achieved by extending the light period by three hours in both the morning and evening (Supplementary Fig. 10). This experimental design revealed a crucial aspect of *phod1* regulation: its expression appears to be induced by light exposure during specific photoinducible phases in the morning and evening. Under short-day conditions, these photoinducible phases likely occur outside the light period, resulting in a lack of bimodal induction of *phod1* expression. However, under long-day conditions, extended light exposure in the morning and evening coincided with these photosensitive phases, leading to the observed bimodal expression pattern of *phod1*. This suggests that *phod1* expression is not simply a response to light itself but rather to light exposure during specific circadian times that are sensitive to photic stimulation. This interpretation aligns with the concept of circadian gating of light responses, in which the circadian clock modulates the sensitivity of the system to light at different times of the day (Bünning, 1936; Pittendrigh and Minis, 1964). scRNA-seq data have shown that *phod1*^+^ cells express *opn5l1c*, a member of the opsin family of photoreceptors (Sato et al., 2018), along with key components of the circadian clock machinery, including *per2a*, *per2b*, *cry1b*, and *hlfa*. In the upstream region of *phod1*, we identified DNA motifs that resemble light-responsive modules (Vatine et al., 2009; Mracek et al., 2012), consisting of an E-box and a D-box (Supplementary Fig. 11). E-box and D-box motifs are known as a key regulatory DNA motif that link the circadian clock function to photoperiodic time measurement (Dardente et al., 2010; Liddle et al., 2022). In addition, light-responsive modules have been shown to play critical roles in light-dependent gene expression, as demonstrated in studies of the *per2* and *cry1a* genes in zebrafish (Vatine et al., 2009; Mracek et al., 2012). The co-expression of *phod1* with photoreceptor and circadian clock genes, and the presence of light-responsive enhancer elements in the upstream region of *phod1* suggests that *phod1*^+^ cells may have the capacity for direct photoreception, potentially allowing cell-autonomous light sensing and circadian entrainment. Further investigation of these factors may provide detailed information of the regulatory mechanisms underlying *phod1* bimodal expression.

Growth hormone plays crucial roles in regulating various aspects of metabolism, including protein, lipid, and carbohydrate metabolism (Møller & Jørgensen, 2009; Vélez and Unniappan, 2021). Through direct actions and stimulation of hepatic insulin-like growth factors (IGFs), the growth hormone exerts significant effects on protein synthesis, lipid metabolism, and carbohydrate metabolism in different target tissues (Møller & Jørgensen, 2009; Vélez and Unniappan, 2021). Interestingly, growth hormone secretion and its downstream effects have been reported to exhibit seasonal patterns in several vertebrates (Fish: Marchant and Peter, 1986; Young et al., 1989; Pérez-Sánchez et al., 1994; Figueroa et al., 2005; Amphibian: Mosconi et al., 1994; Polzonetti-Magni et al., 1995; Bird: John et al., 1983; Mammal: Webster et al., 1996; Blumenthal et al., 2011; Tendler et al., 2021). In addition, the expression and its secretion of the growth hormone in several species have been shown to be regulated by photoperiod (Fish: Björnsson et al., 1989; McCormick et al., 1995; Björnsson, 1997; Björnsson et al., 2000, Mammal: Barentonet al., 1988; Jin et al., 2012). However, the precise mechanisms by which photoperiodic information modulates growth hormone signaling and its metabolic effects remain unknown. In this study, we found that *phod1* is essential for photoperiodic regulation of *gh1* expression. Specifically, while *gh1* expression was increased by a long-day stimulus and decreased by a short-day stimulus in wild-type fish, *phod1* knockout fish showed dysregulated *gh1* expression, irrespective of photoperiodic conditions, suggesting that *phod1* is required for proper photoperiodic control of the growth hormone signaling pathway. scRNA-seq analysis revealed that *phod1*^+^ cells and somatotropes are distinct cell populations in the pituitary, suggesting that *phod1* regulates *gh1* expression through intermediate factors (Supplementary Fig. 12). In addition, *phod1* was predicted to be a membrane protein (Supplementary Fig. 13), indicating that the photoperiod-dependent expression of *phod1* may lead to the secretion of certain factors that influence *gh1* expression in somatotropes. LIANA-based cell-cell communication analysis revealed potential signaling interactions from *phod1*^+^ cells to somatotropes via *inhbaa* and *rln3* (Fig. 3d). Notably, activin A (which consists of a homodimer of inhibin beta A subunits, encoded by *inhbaa*) has been shown to negatively regulate growth hormone synthesis and secretion (Billestrup et al., 1990; Fung et al., 2017). Additionally, rln3 signaling was found to significantly decrease growth hormone levels (Sutton et al., 2009). Although the precise mechanisms remain to be elucidated, cell-cell communication mediated by *phod1* could represent a pathway linking photoperiodic information to growth hormone regulation.

Analysis of *phod1* knockout fish revealed that *phod1* was essential for proper metabolic regulation. In many animals, the transition to spring/summer is accompanied by increased ambient temperatures and food availability, leading to enhanced activity levels as the animals search for food and potential mates. This increased activity is tightly linked to breeding behaviors and requires appropriate metabolic adjustments to meet the higher energy demands (Parker and Cheung, 2020). Indeed, wild-type fish showed increased locomotor activity in response to long-day stimuli, whereas *phod1* knockout fish showed no such photoperiodic response, suggesting a crucial role for *phod1* in the photoperiodic regulation of physical activity. Consistent with this behavioral phenotype, metabolomic analysis revealed significant alterations in energy metabolism in *phod1* knockout fish. We observed accumulation of various metabolites including glycolysis and pentose phosphate pathway intermediates (glucose 6-phosphate, fructose 6-phosphate, and ribulose 5-phosphate), multiple coenzymes and cofactor (NAD, NADPH, FAD, FMN, and CoA), a nucleotide (ADP), nucleosides (guanosine and inosine), and amino acid (arginine) (Bian et al., 2022; Rui, 2014). Particularly noteworthy was the accumulation of coenzymes and metabolic intermediates, suggesting altered cellular energy metabolism. These metabolic changes, along with the decreased locomotor activity, indicate that *phod1* knockout results in significant perturbations in energy utilization. Taken together, we propose that *phod1* may function as a key mediator of seasonal adaptation by modulating not only growth hormone signaling but also cellular energy metabolism in response to increasing day length, thereby helping animals prepare for and sustain the energetically demanding activities of the breeding season.

In conclusion, we demonstrate that *phod1* serves as a key link between photoperiodic sensing and metabolic regulation by modulating of growth hormone signaling. This mechanism enables animals to coordinate their metabolic state with seasonal changes in behavior and energy demands, providing a novel mechanism for seasonal adaptation in vertebrates.

## Methods

### Animals

Medaka (*Oryzias latipes*) were obtained from a local dealer (Fuji 3A Project, Nagoya, Japan). Medaka were kept under short-day (10 h light / 14 h dark; 26°C) conditions or long-day (16 h light / 8 h dark; 26°C) conditions in a housing system (MEITO system, Meito Suien; LP-30LED-8CTAR, NK system). All experiments in this study were performed using adult medaka. All animal studies were carried out in accordance with the ARRIVE guidelines. All methods complied with the relevant guidelines and regulations and were approved by the Animal Experiment Committee of Nagoya University and the National Institutes of Natural Sciences.

### Reverse transcription quantitative PCR

Total RNA (200 ng) was reverse transcribed using the ReverTra Ace qPCR RT Kit (Toyobo). Samples contained SYBR Premix Ex Taq II (Takara), 0.4 μM gene-specific primers (Supplementary Table 6), and 2 µL synthesized cDNA in a 20 µL volume. qPCR was performed in an Applied Biosystems QuantStudio 3 Real-Time PCR System as follows: 95°C for 30 s, followed by 40 cycles of 95°C for 5 s and 60°C for 34 s. *Ribosomal protein L7* (*rpl7*) was used as an internal control.

### In situ hybridization

DNA fragments of 826 bp corresponding to nucleotides 198–1,023 of *LOC101165601* cDNA (XM_004081073.4) were amplified with gene-specific primers (*LOC101165601*: Forward: 5’-TCGCTGGACCTGGGATCTAT-3’; Reverse: 5’-ATTGGAGTCATGACGTCTGGG-3’) and cloned into the pCRII-TOPO verctor using the TOPO TA Cloning Kit (Invitrogen). Following linearization of the oligonucleotide-inserted pCRII-TOPO verctor with EcoRV, digoxigenin (DIG)-labeled cRNA probes were generated using the DIG RNA Labeling Mix and Sp6 RNA polymerase (Roche Diagnostics). In situ hybridization was performed as previously described (Hiraki et al., 2012). In brief, the brain was fixed in 4% paraformaldehyde, embedded in paraffin, and cut into 10 µm sections in the coronal plane. Tissue sections were treated with proteinase K (3 μg/mL) and subsequently hybridized with DIG-labeled RNA probes. Hybridization signals were detected using an alkaline phosphatase-conjugated anti-DIG antibody (Roche Life Science), with Nitro Blue tetrazolium and 5-bromo-4-chloro-3-indolyl-phosphate as chromogens. Color development was allowed to proceed overnight.

### Single-cell RNA-seq analysis

We reanalyzed the single-cell RNA-sequencing (scRNA-seq) data of the medaka (*Oryzias latipes*) pituitary gland published by Siddique et al. (2021). This dataset was obtained from the European Nucleotide Archive (ENA) (accession number: PRJNA683018). Raw sequencing data were processed using Cell Ranger (version 8.0.0, 10x Genomics) to generate gene expression matrices. For read mapping and quantification of gene expression, we used the NCBI RefSeq annotation for the medaka genome (*Oryzias latipes*, ASM223467v1). The filtered count matrices from the four sequencing replicates were combined by adding the same barcodes. The combined counts matrix was further analyzed using the Seurat R package (version 5.0.2) (Hao et al., 2024). Quality control was performed using the DoubletFinder R package (version 2.0.4) (McGinnis et al., 2019), excluding 2% of the total cells as doublets. All parameters for DoubletFinder were determined according to the developer’s recommendations (pN = 0.25, pK = 0.01). Data normalization and feature selection were performed using the SCTransform method. Dropout imputation was performed using ALRA implemented in the SeuratWrappers R package (version 0.3.5). Dimensionality reduction was performed using principal component analysis (PCA), followed by Uniform Manifold Approximation and Projection (UMAP) using the top 20 principal components, which were calculated from the top 3,000 highly variable genes. The cell clustering was performed using the Louvain algorithm with at a resolution of 0.2. The Wilcoxon-rank sum test was performed between clusters to identify characteristic marker genes, and cell types were inferred by comparing their expression patterns with known cell type-specific marker genes. We defined marker genes that were expressed in over 80% of cells in a cluster with fold-change > 5 and an adjusted *P* value < 1.0E-50. All statistical analyses and visualizations were performed using R software (version 4.2.1).

### Cell-cell communication analysis

Cell-cell communication analysis was performed using the LIANA R package (version 0.1.13) (Dimitrov et al., 2022). Ligand-Receptor (L-R) interaction resource provided in LIANA was customized for medaka pituitary gland dataset. L-R pairs in consensus resource were decomplexified, and human gene symbols were converted into medaka ones. To focus on intercellular surface or membrane-bound interactions, only L-R pairs with annotations of “ligand”, “cell_surface_ligand” and “receptor” were retained and constructed custom database consisted of 6,156 medaka L-R gene pairs. LIANA analysis was carried out with this customized database and 5 scoring methods (connectome, logfc, natmi, sca, and cellphonedb). Results from each scoring methods were integrated into an aggregated rank by Robust Rank Aggregation method. L-R pairs with aggregated rank < 0.05, sca L-R score > 0.7, and natmi interaction specificity < 0.15 were defined as significant.

### Genome editing using the CRISPR/Cas9 system

The single guide RNA (sgRNA) expression vector (pDR274; Addgene plasmid 42250) was linearized with BsaI and purified using the NucleoSpin Gel and PCR Clean-up (Takara Bio). Appropriately designed oligonucleotides were synthesized using an oligonucleotide purification cartridge (OPC)-based purification service at Eurofins Genomics. A pair of complementary oligonucleotides was annealed by heating at 95°C for 2 min and then cooled slowly to 25°C for 30 min. The annealed oligonucleotides were ligated into the linearized pDR274 vector. Following linearization of the oligonucleotide-inserted pDR274 vector and the Cas9 expression vector (hCas9; gifted by Prof. Zhang of Massachusetts Institute of Technology) with DraI and NotI respectively, the sgRNAs and the capped Cas9 mRNA were synthesized by using the AmpliScribe T7-Flash Transcription Kit (Epicentre) and the mMESSAGE mMACHINE SP6 Transcription Kit (Thermo Fisher Scientific), respectively. Both RNAs were purified using the RNeasy Mini Kit (QIAGEN). A mixed solution of two sgRNAs (50 ng/µL) and the Cas9 mRNA (100 ng/µL) were microinjected into one-cell stage embryos using a fine glass needle. Genomic DNA was extracted from caudal fin clips of adult fish. Caudal fin clips were lysed individually in 100 µL of Buffer G2 containing 800 mM guanidine-HCl, 30 mM Tris-HCl (pH 8.0), 30 mM EDTA (pH 8.0), 5% Tween-20, and 0.5% Triton X-100 with 1 µL of 10 mg/mL proteinase K (FUJIFILM Wako Pure Chemical Corporation) and incubated at 55°C for 1 h. After centrifugation, 50 µL of the supernatant was purified by ethanol precipitation. Samples were resuspended in 100 µL of Tris-EDTA buffer and used as genomic DNA samples. The DNA region containing *phod1* target site of sgRNA was amplified from the DNA samples using the KAPA Taq Extra PCR Kit (Kapa Biosystems). The PCR primers 5’-AAACAGGCTACTGGGGCAAA-3’ and 5’-TTGACACGTTACTGGTGCTGT-3’ were used for genotyping *phod1* knockout medaka, and amplicons were analyzed by heteroduplex mobility assay using an automated microchip electrophoresis system (MCE-202 MultiNA; Shimazu). Sequences of the mutant alleles were determined by direct Sanger sequencing of the PCR products.

### RNA-seq analysis

For transcriptome analysis of the brain, medaka maintained under short-day conditions were transferred to long-day conditions, and whole brains were collected from three fish per genotype before and 12 days after the transition. The brains were immediately immersed in RNAprotect Tissue Reagent (QIAGEN) for 24 h, and then the region containing the hypothalamus and pituitary was carefully dissected under a microscope. Total RNA was isolated using the RNeasy Micro Kit with DNase I treatment (QIAGEN). Libraries were prepared using the NEBNext Ultra II Directional RNA Library Prep Kit, following poly(A) RNA isolation (New England Biolabs). Paired-end 150 bp sequencing was performed on an Illumina NovaSeq 6000 platform to generate 40 million reads per library (Rhelixa, Inc.). Raw reads were quality-controlled using FastQC (v0.11.9) (Andrews, 2010) and trimmed using the Trimmomatic (v0.39) (Bolger et al., 2014). Clean reads were mapped to the *Oryzias latipes* reference genome using STAR (v2.7.9a) (Dobin et al., 2012). Transcript abundance was quantified as transcripts per million (TPM) using RSEM (v1.3.1) (Li and Dewey, 2011), and differential expression analysis was performed using DESeq2 (v1.38.2) (Love et al., 2014). For the liver transcriptome analysis, samples were collected from five fish per genotype under long-day conditions and immediately frozen. Total RNA was extracted using the Maxwell RSC simplyRNA Tissue Kit with the Maxwell RSC Instrument (Promega). Libraries were prepared using the Lasy-Seq v1.1 protocol for 3’ RNA sequencing (Kamitani et al., 2019; Kashima et al., 2021). Pooled libraries (up to 96 samples) were sequenced on an Illumina HiSeqX Ten platform using paired-end sequencing. Reads were mapped to the *Oryzias latipes* reference genome using BWA (v0.7.17-r1188) (Li, 2013), and transcript abundance was quantified as reads per million mapped reads (RPM) using Salmon (v1.4.0) (Patro et al., 2017). Differential expression analysis was performed using the DESeq2 (v1.44.0).

### LC-MS/MS analysis

Metabolomics analysis was conducted by Shimadzu Techno-Research, Inc. The livers of three wild-type and three knockout medaka maintained under long-day conditions were used. For metabolite extraction, each sample was homogenized in 500 μL of internal standard solution (10 µmol/L 2-morpholinoethanesulfonic acid in methanol) using a Nippi Bio-Masher. After homogenization, 250 μL of water and 500 μL of chloroform were added sequentially and thoroughly mixed. The mixture was centrifuged at 13,000 rpm, 4°C for 15 minutes, after which 400 μL of the upper layer was collected and dried using a centrifugal evaporator. For metabolomic analysis, a Shimadzu LC-30A system coupled with a Shimadzu LCMS-8060 triple quadrupole mass spectrometer was used. Chromatographic separation was achieved on a Mastro2 C18 column (2.0 mm × 150 mm, 3 μm) maintained at 40°C. Mobile phase A consisted of 15 mM acetic acid with 10 mM dipentylamine and mobile phase B consisted of methanol. Gradient elution was employed, starting at 0% B and subsequently increasing to 15% B at 8 min and to 98% B at 12 min, and held until 15 min before returning to the initial conditions. The flow rate was 0.3 mL/min, and the injection volume was 3 μL. Mass spectrometric detection was performed using the LC-MS/MS Method Package for Primary Metabolites Version 3 (Shimadzu Corporation). The peak areas for each compound were normalized to the internal standard and sample weight. Normalized values for the knockout group were compared with those for the wild-type group. Peak areas below 3,000 were considered below the quantification limit and excluded from the analysis.

### Behavioral tests

Each fish was placed in a plastic container (80 mm length × 50 mm width × 50 mm height) and their behavior was tracked for approximately 5 days during the transition from short-day to long-day conditions using an AutoCircaS (TAISEI). The total amount of activity was measured over 10 minutes intervals.

### Statistical analysis

Data are presented as the mean ± SEM. *F*-tests were used to determine the variance. Data with a normal distribution were analyzed by a Student’s *t*-test between two groups. Where the variance was significantly different between groups, a Welch’s *t*-test was used.

## Data availability

The RNA-seq data generated in this study have been deposited in the NCBI’s Gene Expression Omnibus (GEO) (accession number GSE284109, GSE286199). All other data are available from the authors upon request.

## Acknowledgements

We thank the NBRP-Medaka (National Bio-Resource Project of MEXT, Japan) for use of their facilities. Computational resources were provided by the Data Integration and Analysis Facility, National Institute for Basic Biology. We also thank M. Okubo and Y. Ikuno for administrative support; A. Akama, C. Kinoshita, Aki Ieda and A. Matsumiya for fish facility maintenance; T. Shimmura, A. Shinomiya, T. Ohkawa, T.K. Tamai and Y. Saitoh for helpful discussions; H. Nishide and H. Abe for technical assistance. This work was supported by JSPS KAKENHI Grant Number 20K15840 (Grant-in-Aid for Early-Career Scientists), 24K18161 (Grant-in-Aid for Early-Career Scientists), 19H05643 (Grant-in-Aid for Scientific Research (S)), 24H00058 (Grant-in-Aid for Scientific Research (S)) and Narishige Zoological Science Award. WPI-ITbM is supported by the World Premier International Research Center Initiative (WPI), MEXT, Japan.

## Author contributions

T.N. and T.Y. conceived and designed the research. T.N. performed qPCR analysis, in situ hybridization, metabolomic analysis and behavioral assays. T.Ya., R.F. and C.H. performed scRNA-seq analysis. T.N. and M.K. performed RNA-seq analysis. T.N. and M. M. generated *phod1* KO medaka. S.A. and K. N. provided new methods and materials. T.N. and T.Y. wrote the manuscript. All authors discussed the results and commented on the manuscript.

