## Supplementary Figure for "*Photoperiod decoder 1* regulates seasonal changes in energy metabolism through the growth hormone signaling pathway"

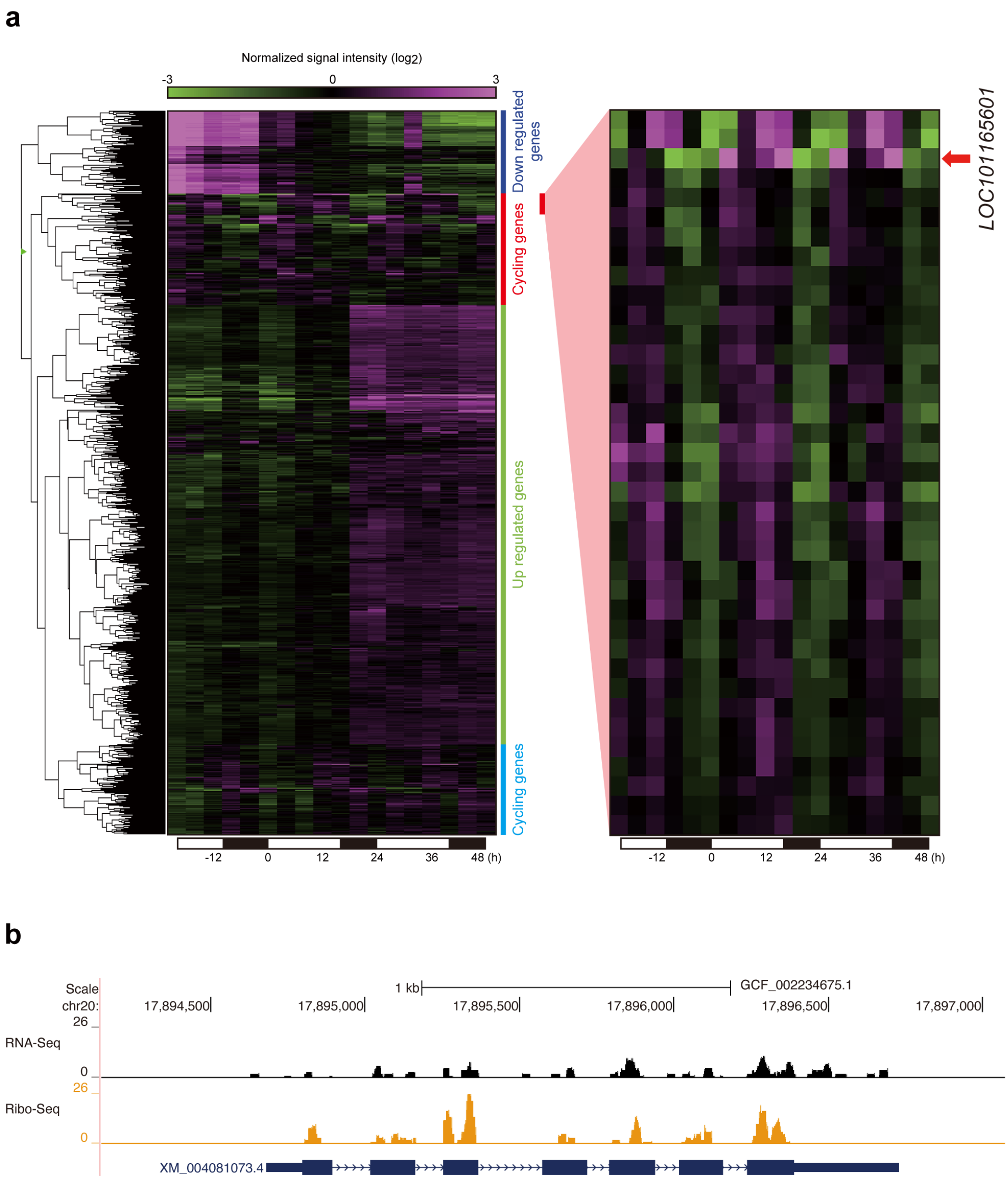


**Supplementary Fig. 1 Identification of uncharacterized gene that shows photoperiodic responsiveness.**

**a** Heatmap of long-day response transcripts (1,249 transcripts) expressed in the brain region containing the hypothalamus and the pituitary, which were identified in a previous study (Nakayama et al., 2019) (left). The data were normalized to the entire data set. The color scale represents the normalized signal intensity. The colored bar on the right represents the clustered categories. Magnification of the red bar region in the cycling genes (right). The red arrow indicates *LOC101165601* expression pattern. **b** Mapping results of RNA-seq and Ribosome profiling (Ribo-seq) analysis at the *LOC101165601* locus (Nakayama et al., 2019). Top panel shows the RNA-seq results (black). The bottom panel shows the Ribo-seq results (orange).

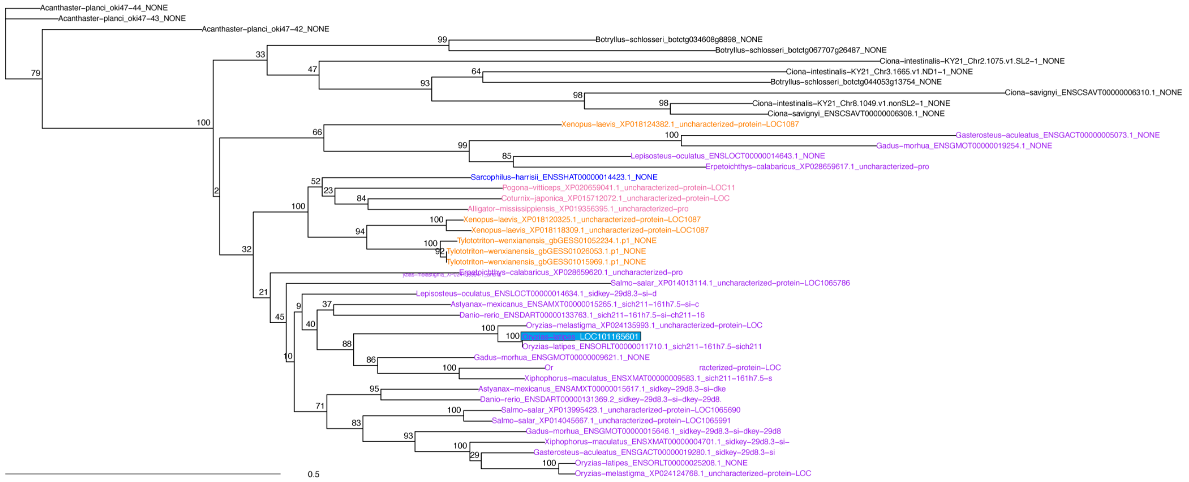


**Supplementary Fig. 2 Phylogenetic distribution of *LOC101165601* in vertebrates.** Phylogenetic tree inferred by ORTHOSCOPE (Inoue and Satoh, 2019) using default parameters (BLASTP with BLOSUM62 matrix, E-value < 0.05). The tree was constructed using Neighbor-joining method with bootstrap analysis (bootstrap values shown at nodes). Scale bar indicates substitutions per site. Query sequence (*LOC101165601* from *Oryzias latipes*) is highlighted in light blue. Other sequences are from the ORTHOSCOPE database, including sequences from fish, amphibians, reptiles, birds, mammals (*Sarcophilus harrisii*), tunicates (*Ciona* and *Botryllus*), and echinoderms (*Acanthaster planci*). Note that ortholog of *LOC101165601* were not found in eutherians.

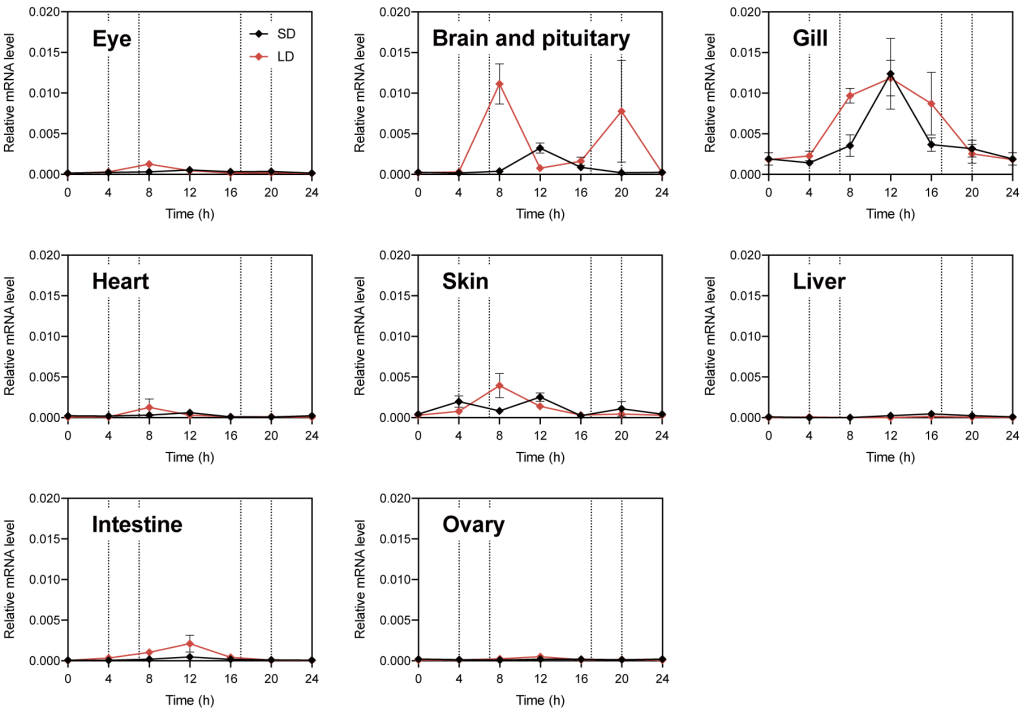


**Supplementary Fig. 3 Expression profile of *LOC101165601* in each tissue under short-day and long-day conditions.**

qPCR was used to measure the relative gene expression levels in each tissue from medaka kept under short-day (10 h light / 14 h dark; 26°C) or long-day conditions (16 h light / 8 h dark; 26°C) (mean ± SEM, and n = 3). Black and red lines indicate expression profiles of *LOC101165601* under short- and long-day conditions, respectively.

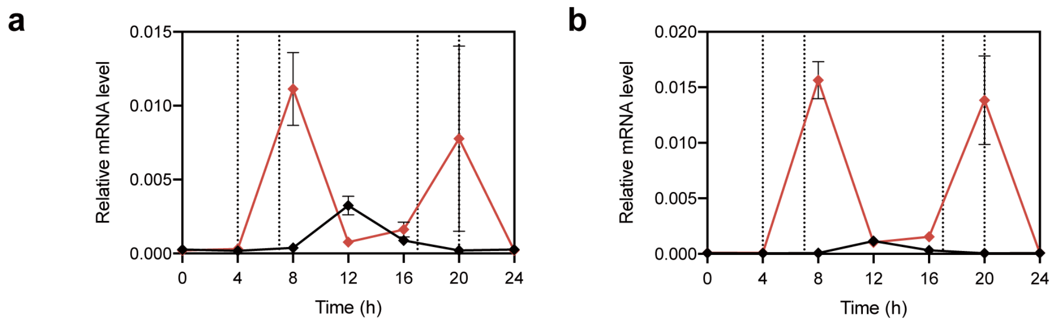


**Supplementary Fig. 4 Expression profile of *LOC101165601* in each sex under short-day and long-day conditions.**

qPCR was used to measure the relative gene expression levels in the brain region containing the hypothalamus and the pituitary of female (a) and male (b) medaka kept under short-day (10 h light / 14 h dark; 26°C) or long-day conditions (16 h light / 8 h dark; 26°C) (mean ± SEM, and n = 3-4). Black and red lines indicate expression profiles of *LOC101165601* under short- and long-day conditions, respectively.

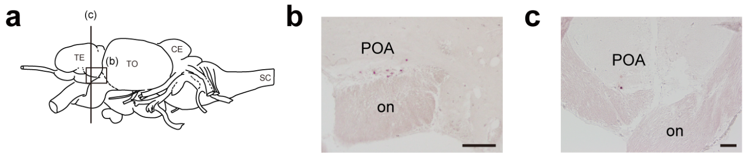


**Supplementary Fig. 5 Expression sites of *phod1* in the preoptic area.**

**a** Schematic illustration of the medaka brain showing the planes of the sections in b and c. TE, telencephalon; TO, tectum opticum; CE, corpus cerebelli; SC, spinal cord. **b** Spatial distribution of *phod1* expression in the preoptic area visualized by in situ hybridization of sagittal brain sections (scale bar = 100 μm). POA, preoptic area; on, optic nerve. **c** Spatial distribution of *phod1* expression in the preoptic area visualized by in situ hybridization of coronal brain sections (scale bar = 50 μm). POA, preoptic area; on, optic nerve.

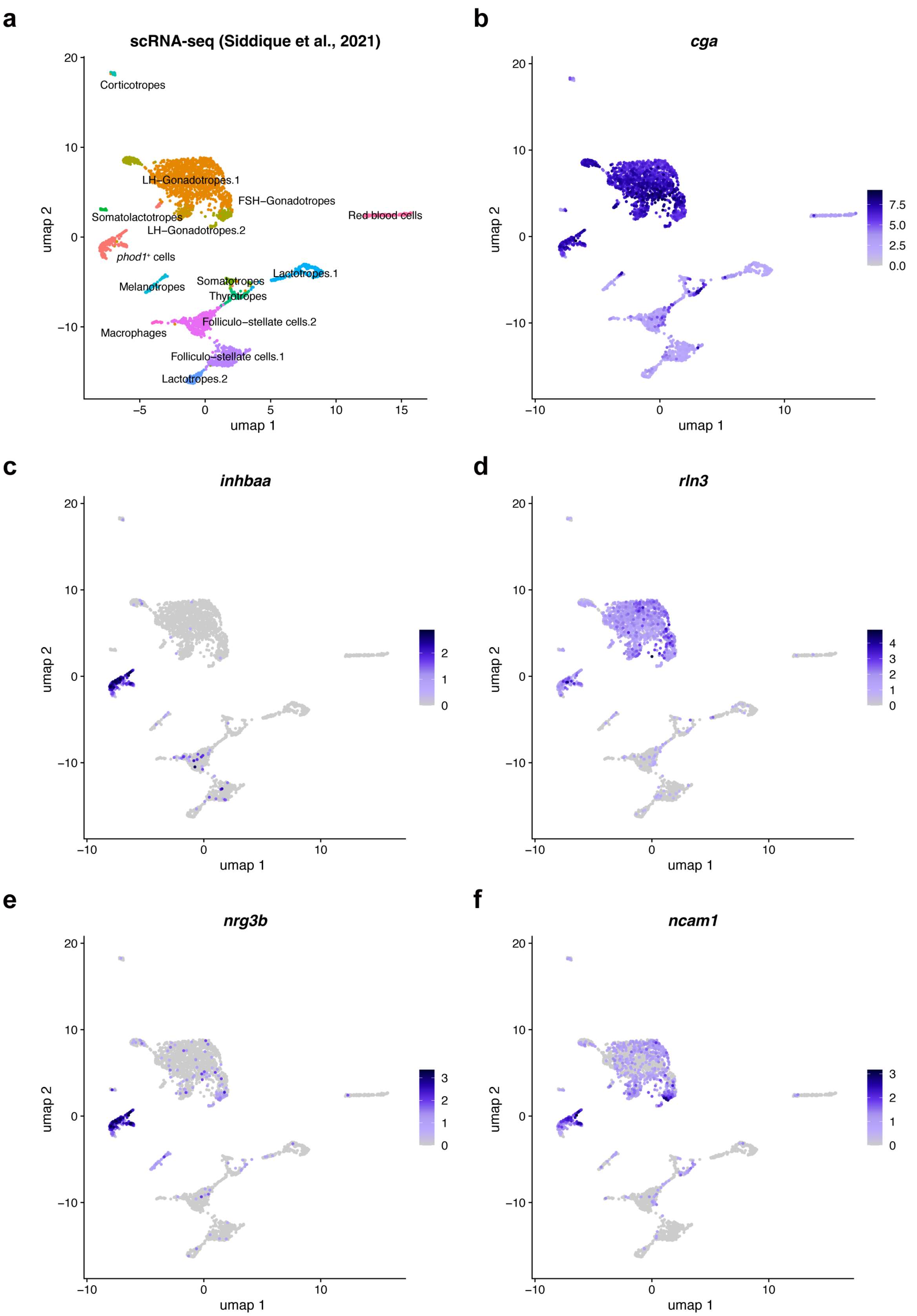


**Supplementary Fig. 6 Single-cell analysis of medaka pituitary.**

**a** Uniform Manifold Approximation and Projection (UMAP) plot of single cell RNA-seq (Siddique et al., 2021). Each dot represents a single cell. **b** Expression of *cga* projected onto the UMAP plot. **c** Expression of *inhbaa* projected onto the UMAP plot. **d** Expression of *rln3* projected onto the UMAP plot. **e** Expression of *nrg3b* projected onto the UMAP plot. **f** Expression of *ncam1* projected onto the UMAP plot.

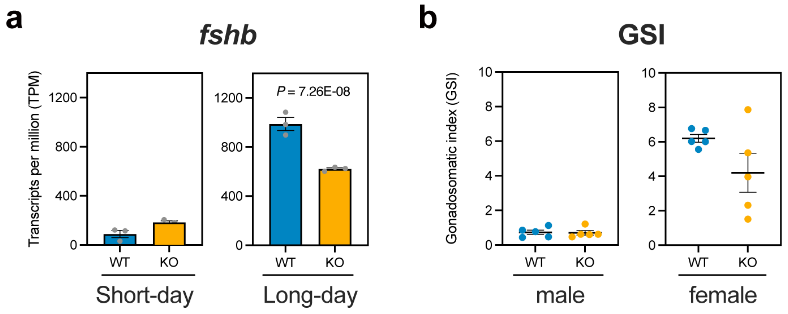


**Supplementary Fig. 7 Expression level of *fshb* and gonadosomatic index (GSI) of wild-type and knockout medaka.**

**a** Expression level of *fshb* in the brain region containing the hypothalamus and the pituitary of wild-type and knockout medaka under short-day and long-day conditions (DESeq2 Wald test, mean ± SEM, and n = 3). **b** GSI of wild-type and knockout medaka under long-day conditions (*t*-test, $P > 0.05$, mean ± SEM, and n = 5). The left and right panels show the GSI of male and female fish, respectively.

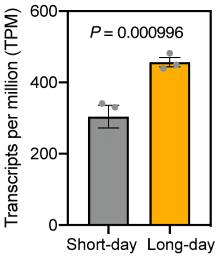


**Supplementary Fig. 8 Expression level of *gh1* in wild-type under short-day and long-day conditions.**

Expression level of *gh1* in the brain region containing the hypothalamus and the pituitary of wild-type medaka under short-day and long-day conditions (DESeq2 Wald test, mean ± SEM, and n = 3). *gh1* expression is induced by long-day stimulation in wild-type.

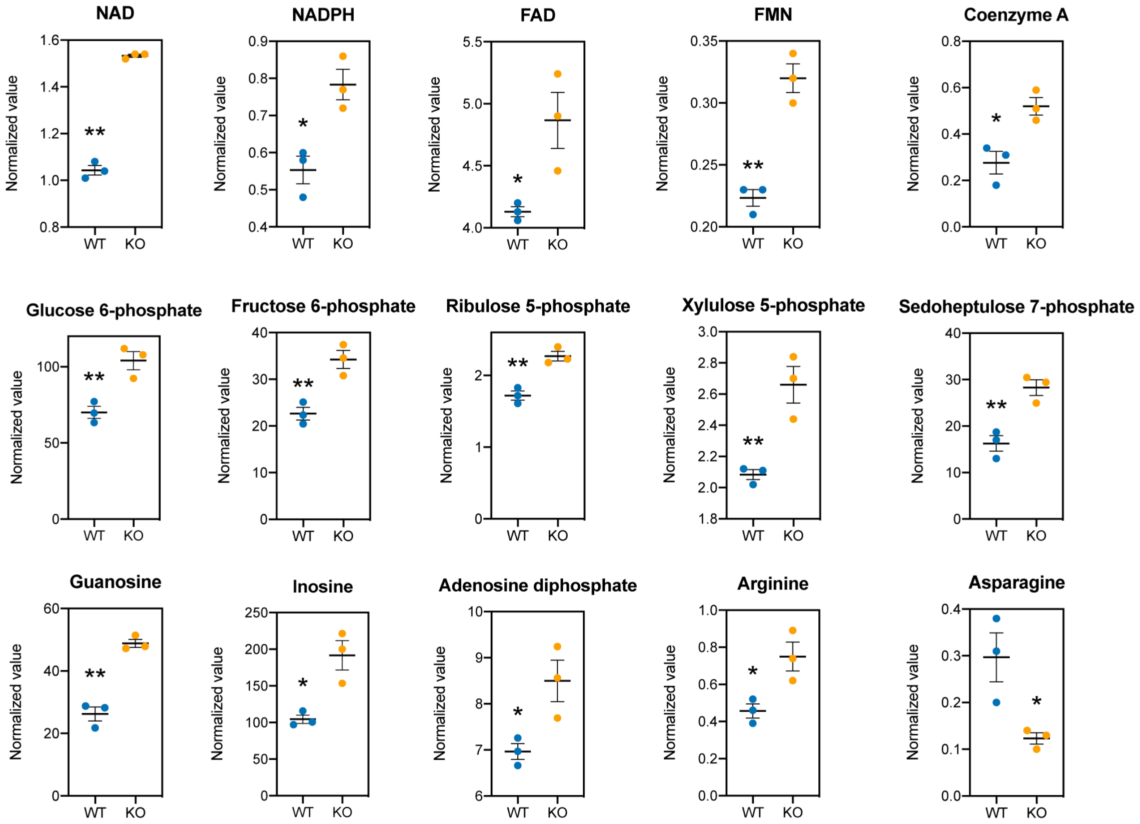


**Supplementary Fig. 9 Metabolomic analyses of the liver.**

Fifteen metabolites with statistically significant differences between wild-type and knockout (Student's *t*-test, **P* < 0.05, ***P* < 0.01, mean ± SEM, and n = 3).

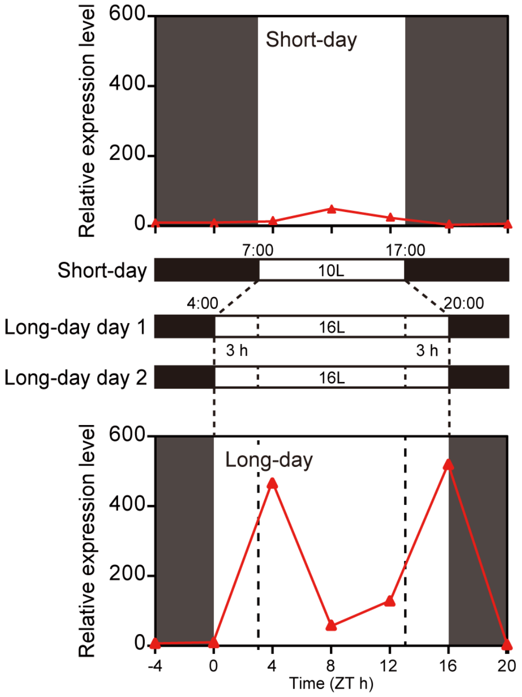


**Supplementary Fig. 10 Schematic drawing of the experimental design.**

Experimental design used identify photoperiodically regulated genes in the previous study (Nakayama et al., 2019). Medaka maintained under short-day conditions (lights on at 7:00, off at 17:00) were transferred to long-day conditions (lights on at 4:00, off at 20:00). A long-day stimulus was achieved by extending the light period by three hours in both the morning and evening

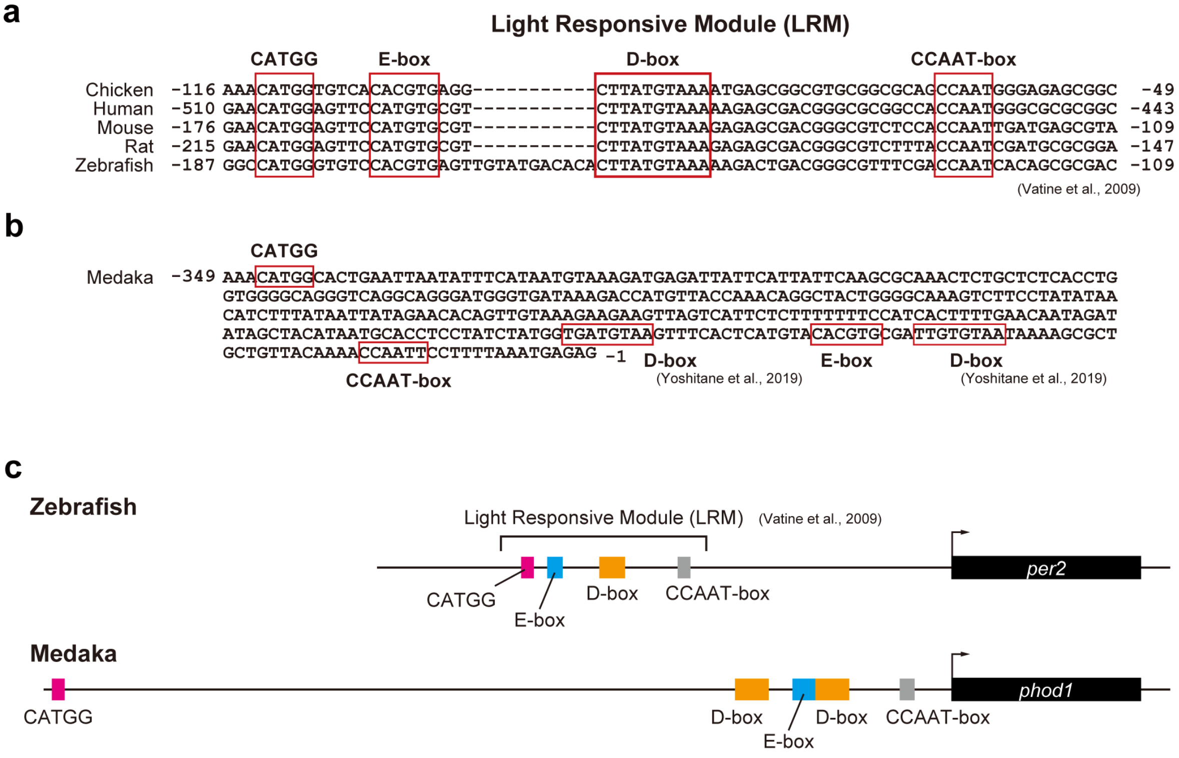


**Supplementary Fig. 11 Light responsive module and DNA motifs in the upstream region of *phod1*.**

**a** Light-responsive module in the upstream region of *per2* in chicken, humans, mice, rats and zebrafish (Vatine et al., 2009). **b** DNA motifs in the upstream region of *phod1* in medaka. D-box elements were identified based on the sequences as described in a previous study (Yoshitane et al., 2019). **c** The light responsive module of *per2* in zebrafish and the upstream region of *phod1* with DNA motifs similar to the light-responsive module.

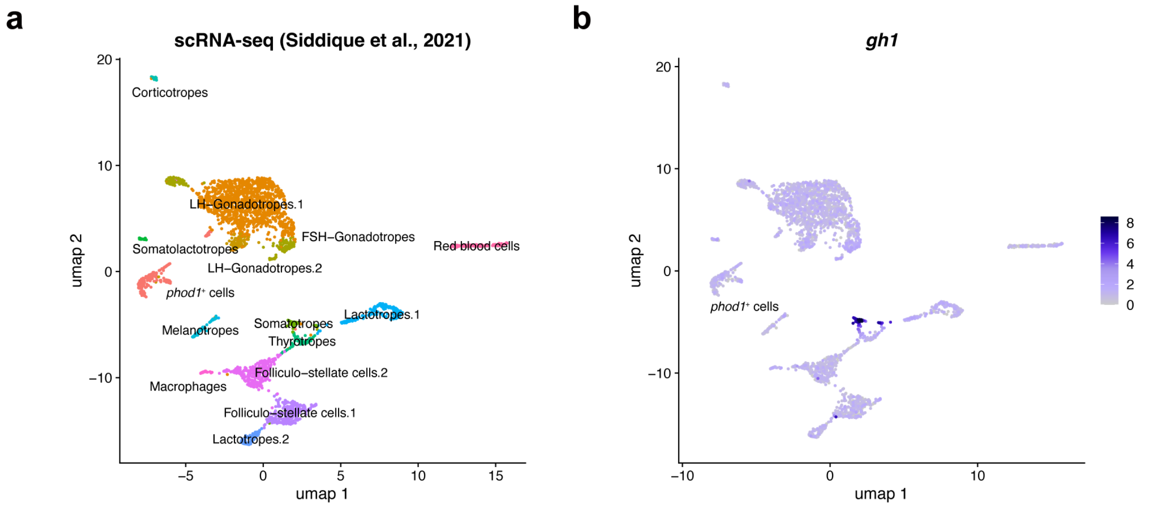


**Supplementary Fig. 12 Single-cell analysis of medaka pituitary.**

**a** Uniform Manifold Approximation and Projection (UMAP) plot of single-cell RNA-seq (Siddique et al., 2021). Each dot represents a single cell. **b** Expression of *gh1* projected onto the UMAP plot.

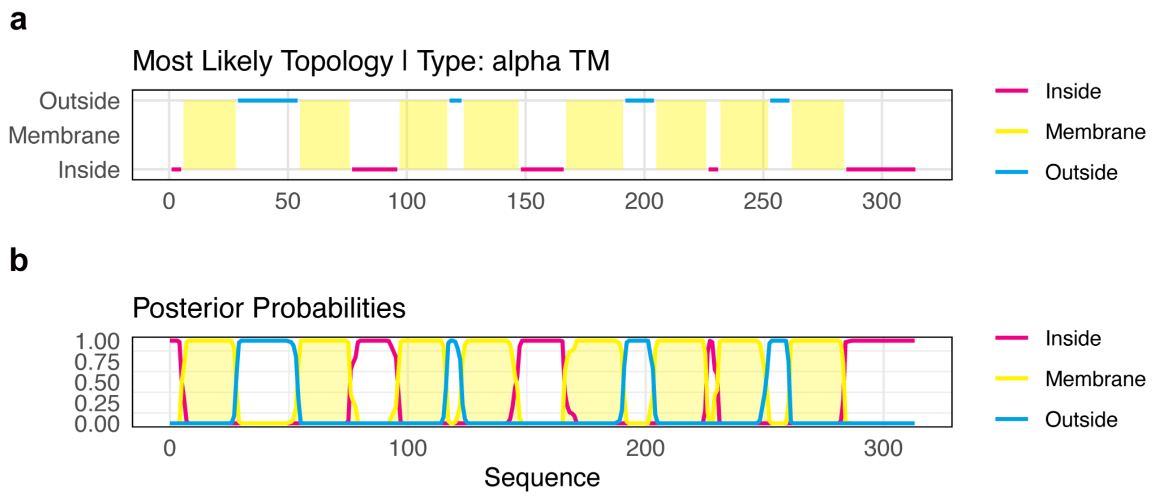


**Supplementary Fig. 13 Transmembrane topology prediction by DeepTMHMM (Hallgren et al., 2022).**

**a** Most Likely Topology prediction showing the predicted transmembrane segments of the protein. The yellow regions indicate the transmembrane segments, whereas pink and light blue regions represent the inside and outside segments, respectively. The analysis predicted eight transmembrane domains characteristic of an alpha-helical transmembrane proteins. **b** Posterior probability plot showing the likelihood of each residue position being in the membrane (yellow), inside (pink), or outside (light blue) environment.
